# Developmental synaptic changes at the transient olivocochlear-inner hair cell synapse

**DOI:** 10.1101/452524

**Authors:** Graciela Kearney, Javier Zorrilla de San Martín, Lucas G. Vattino, Ana Belén Elgoyhen, Carolina Wedemeyer, Eleonora Katz

**Author notes:** **Corresponding Author**: Eleonora Katz, Instituto de Investigaciones en Ingeniería Genética y Biología Molecular “Dr. Héctor N. Torres” (INGEBI-CONICET). Buenos Aires. Argentina, Address: Vuelta de Obligado 2490, 1428 Buenos Aires, Argentina, TE.

## Abstract

In the mature mammalian cochlea, inner hair cells (IHCs) are mainly innervated by afferent fibers that convey sound information to the central nervous system. During postnatal development, however, medial olivocochlear (MOC) efferent fibers transiently innervate the IHCs. The MOC-IHC synapse, functional from postnatal day (P)0 to hearing onset (P12), undergoes dramatic changes in the sensitivity to acetylcholine (ACh) and in the expression of key postsynaptic proteins. To evaluate whether there are associated changes in the properties of ACh release during this period, we used a cochlear preparation from mice at P4, P6-7 and P9-11 and monitored transmitter release from MOC terminals in voltage-clamped IHCs in the whole-cell configuration. The quantum content increased 5.6x from P4 to P9-11 due to increases in the size and replenishment rate of the readily releasable pool (RRP) of synaptic vesicles, without changes in their probability of release (P_vesicle_) or quantum size. This strengthening in transmission was accompanied by changes in the short-term plasticity (STP) properties, which switched from facilitation at P4 to depression at P9-11. We have previously shown that at P9-11, ACh release is supported by P/Q and N-type voltage-gated calcium channels (VGCCs) and negatively regulated by BK potassium channels activated by Ca^2+^ influx through L-type VGCCs. We now show that at P4 and P6-7, release is mediated by P/Q-, R- and L-type VGCCs. Interestingly, L-type VGCCs have a dual role: they both support release and fuel BK channels, suggesting that at immature stages the presynaptic proteins involved in release are less compartmentalized.

**Significance statement:** During postnatal development prior to the onset of hearing, cochlear IHCs present spontaneous Ca^2+^ action potentials which release glutamate at the first auditory synapse in the absence of sound stimulation. The IHC Ca^2+^ action potential frequency pattern, which is crucial for the correct establishment and function of the auditory system, is regulated by the efferent MOC system that transiently innervates IHCs during this period. We show short-term synaptic plasticity properties of the MOC-IHC synapse that tightly shape this critical developmental period.

## Introduction

In the mammalian cochlea, sounds are converted into electrical signals by mechanosensory cells: inner and outer hair cells, IHCs and OHCs, respectively. IHCs convey these signals to the central nervous system, whereas OHCs are involved in the amplification and fine tuning of sounds (Hudspeth, 1997). Sound processing in the cochlea is modulated by descending efferent fibers of the medial olivocochlear system (MOC) (Guinan, 2011). In the mature mammalian cochlea, IHCs are mainly innervated by afferent fibers. However, during postnatal development, before the onset of hearing (postnatal day (P) 12 in altricial rodents), a transient efferent innervation is found in the IHCs (Glowatzki and Fuchs, 2000, Katz *et al.*, 2004, Roux *et al.*, 2011) even before MOC fibers contact their final targets, the OHCs (Liberman *et al.*, 1990, Simmons *et al.*, 1996, Simmons, 2002). Several studies suggest that this transient efferent innervation plays a role in the ultimate functional maturation of cochlear hair cells (Simmons, 2002) and in the correct establishment of the auditory pathway, by regulating the spontaneous IHC firing frequency during this critical developmental period (Glowatzki and Fuchs, 2000, Goutman *et al.*, 2005, Johnson *et al.,* 2013, Sendin *et al.,* 2014, Moglie *et al.,* 2018, Wedemeyer *et al.,* 2018). This notion is reinforced by the observation that mice that lack functional efferent MOC-hair cell synapses present a decreased tonotopic organization in the lateral superior olive and a concomitant impairment in the detection of sound frequency changes (Clause *et al.,* 2014, Clause *et al.,* 2017).

The postsynaptic events at MOC-hair cell synapses are well characterized, namely the activation of calcium permeable α9α10 nicotinic acetylcholine (ACh) receptors (nAChRs) (Elgoyhen *et al.,* 1994, Elgoyhen *et al.,* 2001, Elgoyhen and Katz, 2012) leads to the opening of calcium-dependent K^+^-channels that hyperpolarize hair cells (Dulon and Lenoir, 1996, Glowatzki and Fuchs, 2000, Oliver *et al.,* 2000, Katz *et al.,* 2004, Wersinger *et al.,* 2010, Katz *et al.,* 2011, Wersinger and Fuchs, 2011). In mammals, fast synaptic transmission at both central and peripheral synapses is mediated by multiple types of voltage-gated calcium channels (VGCCs), including N-, P/Q- and R-type (Katz *et al.,* 1997, Plant *et al.,* 1998, Reid *et al.,* 2003, Catterall and Few, 2008). VGCC are formed by at least four different subunits (α1, α2-δ, β, sometimes also γ). The biophysical and pharmacological diversity of VGCC which have been classified into L-, P/Q-, N-, R and T-type arises from the existence of multiple pore-forming α1 subunits (Catterall, 1998, Catterall and Few, 2008). In the mouse cochlea, expression of the α1 subunits Ca_v_1.2 (L-type), Ca_v_1.3 (L-type) and Ca_v_2.3 (R-type) has been shown by PCR analysis (Green *et al.,* 1996).

The MOC-IHC synapse undergoes dramatic changes in innervation pattern, sensitivity to ACh and expression of key postsynaptic proteins (Glowatzki and Fuchs, 2000, Simmons, 2002, Katz *et al.,* 2004, Roux *et al.,* 2011). In developing synapses, synaptic modifications likely take place concurrently in both postsynaptic cells and presynaptic terminals. Thus, the subtypes of Ca^2+^ channels coupled to the release process are developmentally regulated both at central synapses (Iwasaki *et al.,* 2000, Momiyama, 2003, Fedchyshyn and Wang, 2005) and at the neuromuscular junction (NMJ) (Rosato Siri and Uchitel, 1999). We have previously shown that at MOC-IHC synapses from P9-11 mice, ACh release is supported by both P/Q and N-type VGCC and negatively regulated by BK potassium channels activated by Ca^2^+ entry through L-type VGCC (Zorrilla de San Martín *et al.,* 2010). In this work we investigated presynaptic changes that shape the maturation of the MOC-IHC synapse, by analyzing the strength of ACh release, the STP pattern and the types of VGCC coupled to the release process at three postnatal ages P4, P6-7 and P9-11. We describe the developmental changes that take place to provide a tight control of synaptic transmission at the transient efferent synapse. This is consistent with the crucial role played by the MOC-IHC synapse in the establishment of the auditory pathway before hearing onset.

## Methods

### Animal procedures and isolation of the organ of Corti

Procedures for preparing and recording from the postnatal mouse organ of Corti were essentially identical to those published previously (Zorrilla de San Martín *et al.,* 2010). Briefly, apical turns of the organ of Corti were excised from BalbC mice at three postnatal stages: P4, P6-7 and P9-11 (day of birth was considered P0), and used within 3 hours. Cochlear preparations were placed in the chamber for electrophysiological recordings, mounted under a Leica DMLFS microscope and viewed with differential interference contrast (DIC) using a 40x water immersion objective and a camera with contrast enhancement (Hamamatsu C7500-50). All experimental protocols were performed in accordance with the *American Veterinary Medical Associations’ AVMA Guidelines on Euthanasia* (2013 Edition).

### Electrophysiological recordings

IHCs were identified visually and by the size of their capacitance (7-12 pF). The cochlear preparation was continuously superfused by means of a peristaltic pump (Gilson Minipulse 3, with 8 channels, Bioesanco, Buenos Aires, Argentina) containing an extracellular saline solution of an ionic composition similar to that of the perilymph (mM): 144 NaCl, 5.8 KCl, 1.3 CaCl_2_, 0.7 NaH_2_PO_4_, 5.6 D-glucose, and 10 Hepes buffer; pH 7.4. Working solutions containing the different drugs and toxins used were made up in this same saline and delivered through the perfusion system. The pipette solution was (in mM): 140 KCl, 3.5 MgCl_2_, 0.1 CaCl_2_, glycol-bis(2-aminoethylether)-N, N,N', N'-tetraacetic acid (5 mM EGTA), 5 Hepes buffer, 2.5 Na_2_ATP, pH 7.2. Some cells were removed to access IHCs, but mostly the pipette moved through the tissue using positive fluid flow to clear the tip. Currents in IHCs were recorded in the whole-cell patch-clamp mode using an Axopatch 200B amplifier, low-pass filtered at 2-10 kHz and digitized at 5-20 kHz with a Digidata 1322A board (Molecular Devices, Sunnyvale, CA, USA). Recordings were made at room temperature (22-25 °C). Glass pipettes, 1.2 mm i.d., had resistances of 4-7 MΩ. Indicated holding potentials were not corrected for liquid junction potentials (−4 mV). Unless otherwise stated, IHCs were voltage-clamped at a holding voltage of −90 mV. As control of IHC viability before studying transmitter release at the MOC-IHC synapse, once in the whole-cell configuration just after break-in, voltage-protocols were applied to check for the IHCs characteristic voltage-dependent currents (Kros *et al.,* 1998).

### Electrically-evoked transmitter release

Neurotransmitter release was evoked by electrical stimulation of MOC efferent axons. Briefly, the electrical stimulus was delivered via a glass pipette (0.4 to 0.9 MΩ resistance) placed at around 80-100 μm modiolar to the base of the IHC under study. The position of the pipette was adjusted until post-synaptic currents in the IHC were consistently activated. A Grass Stimulator (model S48 stimulator coupled to a SIU5 isolation unit) was triggered via the data-acquisition computer to generate pulses up to 70-300 μA, 0.1-0.2 ms.

### Transmitter release evoked by high extracellular K^+^

Neurotransmitter release from efferent terminals was elicited by depolarization using 15-25 mM external potassium saline applied by a bath perfusion system (rate flow 2 ml/min). Spontaneous synaptic currents (sIPSC) were identified by eye using MiniAnalysis (RRID:SCR_002184). After incubation of the preparation in high K^+^ (15-25 mM) solution for 4-8 min, either an L-type VGCC antagonist (Nifedipine 3 μM) or an agonist (Bay-K 10 μM) was applied through the bath-perfusion system. To quantify the effects of each drug, sIPSC frequency was determined after 15 min of incubation with either Nifedipine or Bay-K and compared to sIPSC frequency in high K^+^ solution without drugs.

### Estimation of the quantum content of transmitter release

The quantum content of transmitter release (*m*) was estimated as the ratio between the mean amplitude of evoked synaptic currents (eIPSC) and the mean amplitude of spontaneous synaptic currents (sIPSC) and by the failures method (del Castillo & Katz, 1954). In order to estimate eIPSC mean amplitude, protocols of 100 stimuli were applied at a frequency of 1 Hz. Spontaneous synaptic currents were recorded during the stimulation protocol. To quantify the effects of each drug or toxin, the preparation was incubated for the time necessary to reach a plateau in the observed effect. Percentage quantum content (% *m*) was calculated as: *m*_t_/*m*_c_*100; where *m*_c_ is the estimation of *m* in the control condition and *m*_t_ is the quantum content estimated after incubation of the preparation with the drug or toxin under study. Synaptic currents were analyzed with MiniAnalysis (RRID:SCR_002184) and with custom routines implemented in Igor Pro 6.0 (Wavemetrics, Igor Pro, RRID:SCR_000325).

### Estimation of short-term synaptic plasticity

The short term plasticity (STP) pattern was studied applying 10-80 repetitions of 10-pulse stimulation trains at 10, 40, and 100 Hz, at 15 (for 10 Hz trains) or 20 s intervals (for 40 and 100 Hz trains). For every pulse, the current amplitude was computed as the difference between the peak of the response and the baseline, which was considered as the current value before the pulse. The probability of successfully evoking a release event (P_success_) for each pulse was computed as the ratio between the number of eIPSCs and the number of sweeps. The average amplitude (A) was obtained by averaging the successful eIPSCs amplitudes after each pulse. The overall amplitude (S) for each stimulus was computed as the average of all recorded amplitudes, including failures of response. To establish the extent of facilitation or depression during a train, the values for the parameters S, P_success_, and A for each pulse were compared to the value at the first pulse.

### Estimation of the readily releasable pool of vesicles (RRP) parameters

The RRP size and the replenishment rate were estimated by the method established by Schneggenburger et al., 1999 (Schneggenburger *et al.,* 1999), based on the use of high frequency stimulation trains to exhaust the synapse and reach a stationary state were only newly recruited vesicles are released. This method assumes that replenishment is constant during the train; it does not assume that all vesicles have the same initial probability and does not require that it remains constant during the train (Schneggenburger *et al.,* 1999, Neher, 2015).

Train stimulation protocols composed of 50 pulses at 100 Hz were repeated 10-30 times at 20 s intervals. The cumulative amplitude mean was plotted as a function of stimulus number and the stationary region was fitted to a linear function. The accumulated amplitude of eIPSCs at the i^th^ pulse was calculated as follows:

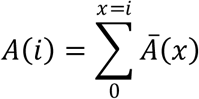

Where *Ā*(*x*) is the mean amplitude of the IPSCs evoked by the x^th^ pulse of the train. To guarantee that the stationary region of the cumulative plot had been considered, the linear fit was calculated on the accumulated amplitudes of the last 20 pulses of the train. In this region, the plots of accumulated amplitudes of all recorded cells showed a linear behavior. The intercept of this fit, divided by the quantum size (the mean sIPSC amplitude) corresponds to the number of vesicles that were ready to be released at the beginning of the stimulation train. The slope of the same fit, divided by the quantum size and multiplied by the stimulation frequency corresponds to the replenishment rate expressed as number of vesicles/second. By calculating the ratio between the average number of vesicles released per action potential (quantum content) and the number of vesicles ready to be released (RRP size), the probability of each vesicle to be released (P_vesicle_) was obtained for each cell. For an extensive discussion of the method, its advantages and limitations and comparison with other methods see Neher 2015 (Neher, 2015).

### Statistics

All statistical tests were performed with RStudio 1.1.442 (RRID:SCR_000432) (Team, 2008), except for two-way ANOVA, which was performed with Graphpad Prism 6.01 (RRID:SCR_002798). Before performing any analysis, data were tested for normal distribution using the Shapiro-Wilk normality test, and parametric or nonparametric tests were applied accordingly. For statistical analyses with two datasets, two-tailed paired t test or Wilcoxon signed-rank test were used. For comparisons of more than two paired datasets, one-way repeated measures ANOVA followed by Tukey’s post-hoc test, or Friedman test followed by Conover’s post-hoc test or two-way repeated measures ANOVA followed by Bonferroni’s post-hoc test were applied. For comparisons of more than two unpaired datasets, one-way ANOVA followed by Tukey’s post-hoc test or Kruskal–Wallis test followed by Dunn’s post-hoc test were used. In these cases, the *p* value reported corresponds to the *p* value of the post-hoc test, unless otherwise stated. Values of *p* <0.05 were considered significant. All data were expressed as mean ± S.E.M., unless otherwise stated.

### Drugs and Toxins

Stock solutions of dihydropyridines (Nifedipine and (±)-BayK-8644, Bay-K) were prepared in dimethyl sulfoxide (final concentration ≤0.1%). Peptidic toxin stock solutions were prepared in distilled water. Reagents and Iberiotoxin were purchased from Sigma-Aldrich Co. (St. Louis, MO, USA), Iberiotoxin was purchased also from Tocris Bioscience (Bristol, UK), all other toxins and drugs were purchased from Alomone Labs (Jerusalem, Israel). All drugs and toxins were thawed and diluted in the extracellular solution just prior to use.

## Results

### Properties of the MOC-IHC synaptic transmission during postnatal development

#### Synaptic strength increases from P4 to P11

From birth to hearing onset (P12), the MOC-IHC synapse undergoes dramatic changes in the sensitivity to ACh and in the expression of key postsynaptic proteins (Katz *et al.,* 2004, Roux *et al.,* 2011). To investigate whether these postsynaptic developmental changes were matched by presynaptic modifications, we first studied the strength of transmitter release at three postnatal (P) stages (P4, P6-7 and P9-11). To this end we evaluated the quantum content of electrically evoked ACh release and P_success_ at these stages. Both parameters significantly increased from P4 to P11 (Figure 1 b, c and d). The quantum content evaluated by the direct method (see Materials and Methods) was: P4 = 0.20 ± 0.02, n = 18 cells, 17 mice; P6-7 = 0.58 ± 0.06, n = 24 cells, 24 mice; P9-11 = 1.12 ± 0.11, n = 24 cells, 24 mice (P6-7 vs P4 *p* = 0.0036; P6-7 vs P9-11 *p* = 7.40e^−06^; P4 vs P9-11 *p* = 8.33e^−04^). The quantum content values obtained by the failures method in the same cells (see Materials and Methods) were not significantly different to those obtained by the direct method (P4 = 0.20 ± 0.02; P6-7 = 0.54 ± 0.06; P9-11 = 1.16 ± 0.11). Psuccess was: P4 = 0.18 ± 0.02; P6-7 = 0.40 ± 0.04; P9-11 = 0.64 ± 0.04 (P6-7 vs P4 *p* = 3.22e^−05^; P6-7 vs P9-11 *p* = 2.20e^−06^; P4 vs P9-11 *p* = 1.20e^−06^). This shows that synaptic transmission is significantly strengthened along this early postnatal developmental period.

**Figure 1.**
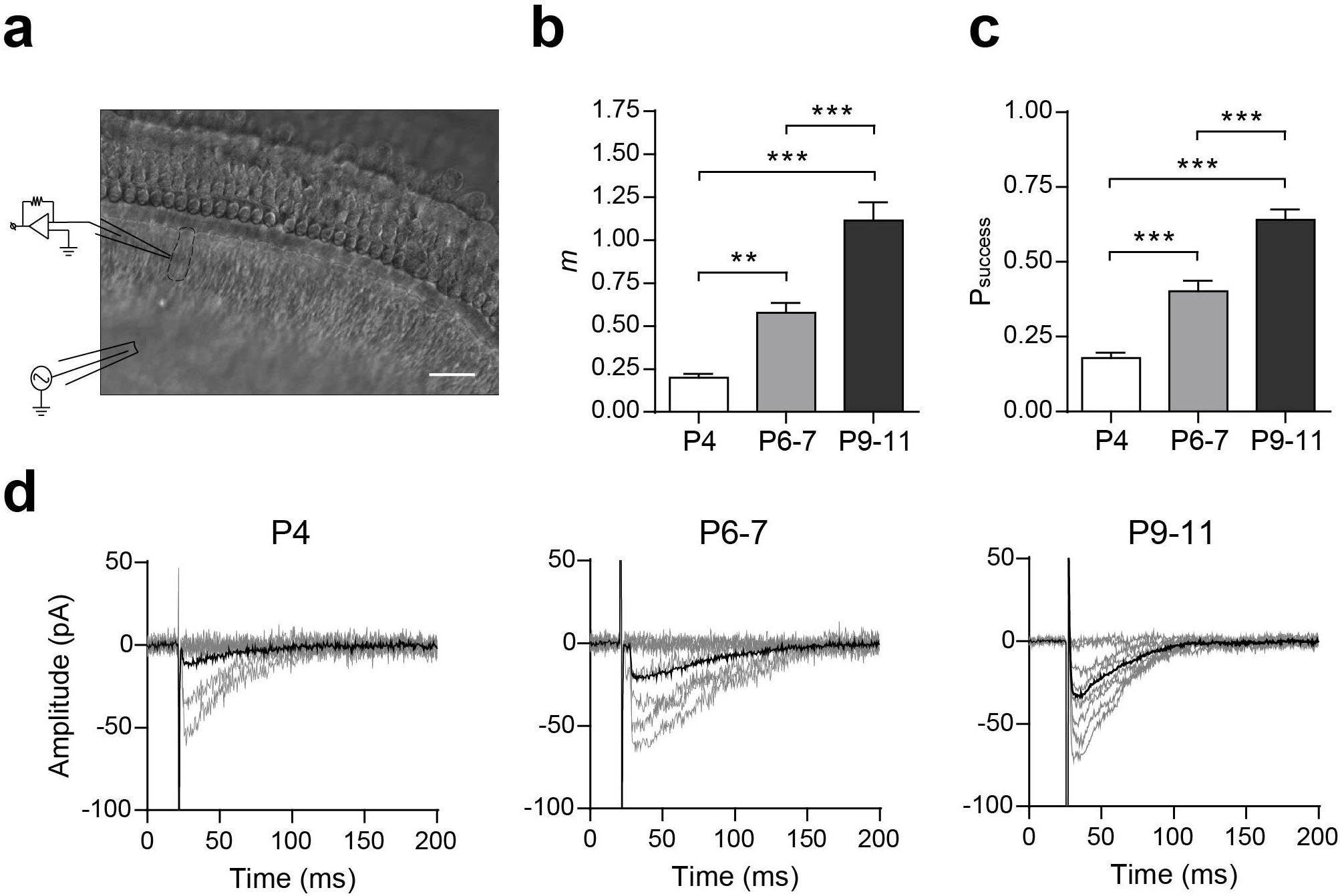
Synaptic strength at the mouse MOC-IHC synapse increases during postnatal development. a, Micrograph of the mouse cochlear preparation used in the present study. IHCs were recorded with a patch pipette at a holding potential of −90 mV while MOC fibers were electrically stimulated using a monopolar electrode placed ∼ 80 μm modiolar to the base of the IHCs. Scale bar, 20 μm. b, Bar graph showing the progressive increase in *m* from P4 to P9-11. c, Bar graph illustrating the increase in the probability of release (P_success_) during the same period as shown in b. d, Representative individual (gray) and average (black) traces of elPSCs recorded at P4, P6-7 and P9-11 IHCs. Data from P911 are shown for comparison and were taken from Zorrilla de San Martín et al., 2010. Error bars are S.E.M. \*\**p*<0.01, \*\*\**p*<0.001.

We analyzed the amplitude and kinetics of the spontaneous postsynaptic currents (sIPSC, Figure 2). No significant changes either in sIPSC amplitude (P4 = 21.16 ± 1.47 pA, n = 20 cells, 19 mice, 550 events; P6-7= 20.06 ± 0.81 pA, n = 20 cells, 20 mice, 462 events; P9-11 = 20.62 ± 1.34 pA, n = 20 cells, 18 mice, 585 events; *p* = 0.9755; Figure 2 b, c) or kinetics (decay time constant P4 = 30.90 ± 1.62 ms, P6-7 = 34.50 ± 2.39 ms, P9-11 = 31.58 ± 1.95 ms; same cells and events as those used for amplitude measurements; *p* = 0.41; Figure 2 b-d) were observed between the different stages. The fact that no changes were observed in sIPCs amplitudes indicates that the quantum size remains constant across the different ages studied.

**Figure 2.**
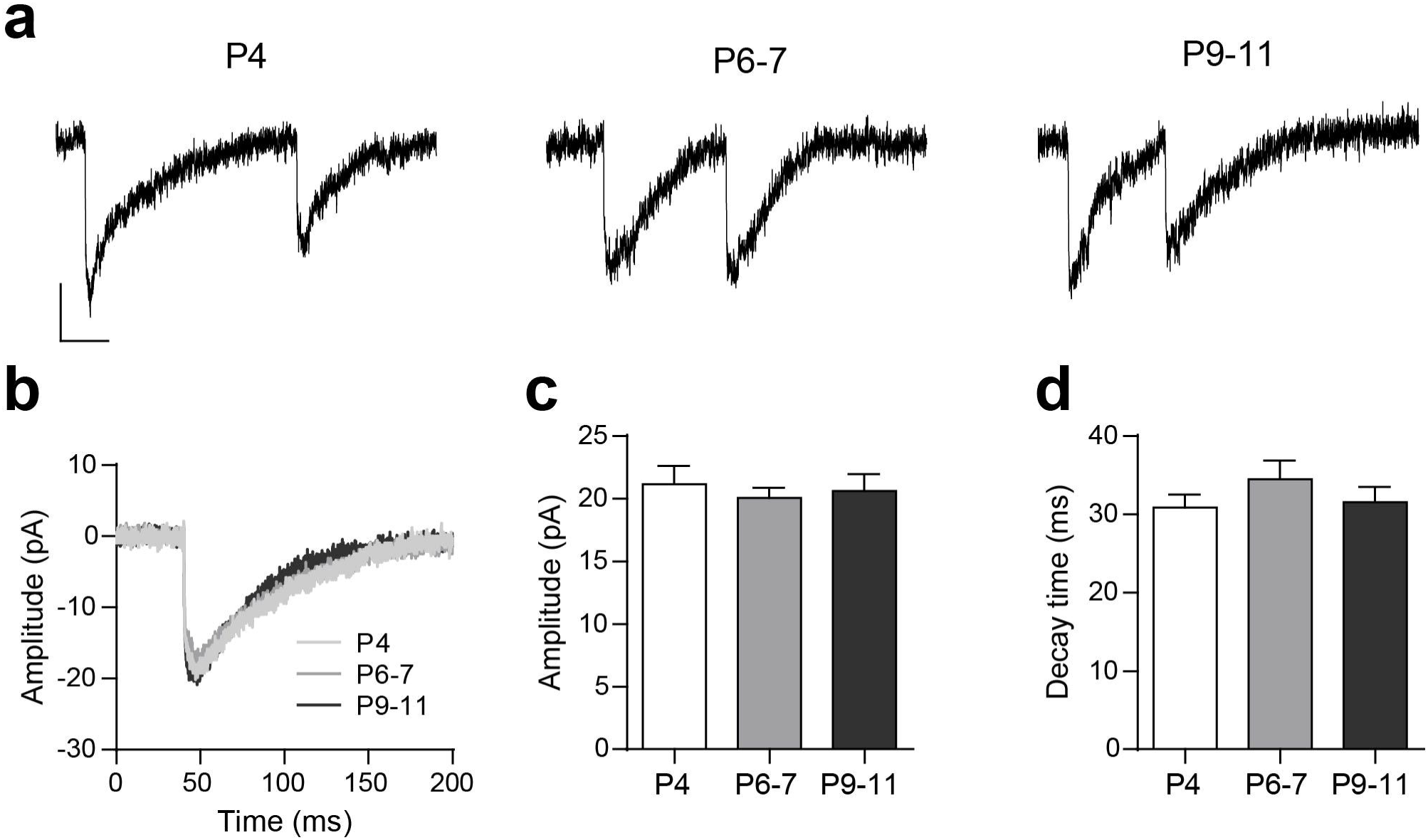
Biophysical properties of sIPSCs at the MOC-IHC synapse during postnatal development. a, Representative traces of sIPSCs recorded at P4, P6-7 and P9-11 IHCs (holding potential: −90 mV). Vertical scale bar: 10 pA, horizontal scale bar: 50 ms. b, Representative average traces of sIPSCs recorded in one IHC from each developmental stage (sIPSCs, n: P4 = 23, P6-7 = 18, P9-11 = 27). c and d, Bar graphs illustrating that neither the sIPSC amplitude (c) nor the decay time constant (d) are modified during MOC-IHC synapse maturation. Error bars are S.E.M.

We also analyzed the frequency of occurrence of sIPSPs and found that at P4, from 4 cells observed during 5-15 min, 3 were completely silent and only 1 cell had a sIPSC frequency of 0.01Hz. At P6-7, sIPSC frequency was 0.02 ± 0.01 Hz, n = 17 cells, 17 mice and it significantly increased at P9-11: 0.18 ± 0.07, n = 10 cells, 8 mice, *p* = 0.0095 (data not illustrated). The increase in the frequency of spontaneous events is consistent with the increment observed in the quantum content of evoked release during the three developmental stages analyzed.

#### Developmental changes in short-term synaptic plasticity

Synapses are endowed with an extraordinary capacity to change according to their previous history. This gives rise to several forms of activity dependent synaptic plasticity that shape synaptic output (Zucker and Regehr, 2002). Synapses with a high initial probability of release tend to depress, whereas those with a low initial probability of release tend to facilitate when challenged by closely spaced stimuli (Fioravante and Regehr, 2011). Facilitation upon high frequency stimulation has been observed for efferent synapses on turtle auditory cells (Art *et al.,* 1984), neonatal rat MOC-IHC synapses (Goutman *et al.,* 2005) and also in mouse MOC-OHC synapses at the onset of hearing (Ballestero *et al.,* 2011). In view of the significant differences in synaptic strength observed at the MOC-IHC synapse between P4, P6-7 and P9-11, we studied whether these differences were reflected in their STP pattern. To this end, we studied the behavior of the MOC-IHC synapse upon application of high frequency stimulation trains. We analyzed the overall response (S), the probability of successfully evoking a release event (Psuccess) and the average amplitude of successful responses (A) along high frequency (10, 40 and 100 Hz) stimulation 10-pulse trains. The normalized values along the train are shown in Figure 3 (S in upper panels, P_success_ in middle panels, A in lower panels). Normalized values > 1 indicate facilitation, while values < 1 indicate depression (Katz and Miledi, 1968, Goutman *et al.,* 2005, Ballestero *et al.,* 2011). Synapses from P4 mice tend to facilitate at the three frequencies tested and along the whole 10-pulse protocols. At all frequencies, the increase in S is accounted for by a reduction in the number of failures and a concomitant increase in P_success_ with no significant changes in A. It must be noted that at 100 Hz, for the maximum facilitation value, the statistical analysis indicates that neither S nor P_success_ were significantly different from 1 (see Table 1). This lack of statistical significance might be due to the high variability in the responses at this early stage. Synapses from P6-7 mice presented significant facilitation at the three frequencies tested, reaching a maximum at the 4^th^ pulse of the 10-pulse protocol and then faded away (Figure 3, Table 1). This suggests that during the train the mechanisms leading to depression balance those leading to facilitation. At this stage, the increase in S is also accounted for by an increase in P_success_ and at 100 Hz, there was also a slight but significant increment in A. This increment in A suggests there might be an increase in the average number of vesicles released during the first part of the 100 Hz train. At P9-11, there was a significant reduction in S and P_success_ at 10, 40 and 100 Hz, being more pronounced at 100 Hz towards the end of the stimulation train (Table 1, Figure 3). A significant reduction in A was also found at 100 Hz, suggesting either a reduction in the average number of vesicles released along the high-frequency stimulation train or a post-synaptic effect due to saturation or desensitization of the postsynaptic receptors (Scheuss *et al.,* 2002, Taschenberger *et al.,* 2005). However, receptor saturation and/or desensitization of the IHC α9α10 nAChR under these conditions are not likely as the exogenous application of 1 mM ACh to cochlear IHCs can evoke currents >0.5 nA with desensitization time constants significantly longer than the duration of the stimulation trains used in the present work (τ decay = 13.8 ± 0.5 s and τ decay = 37.9 ± 4.2 s for the continuous and intermittent application of ACh, respectively (Gomez-Casati *et al.,* 2005).

**Figure 3.**
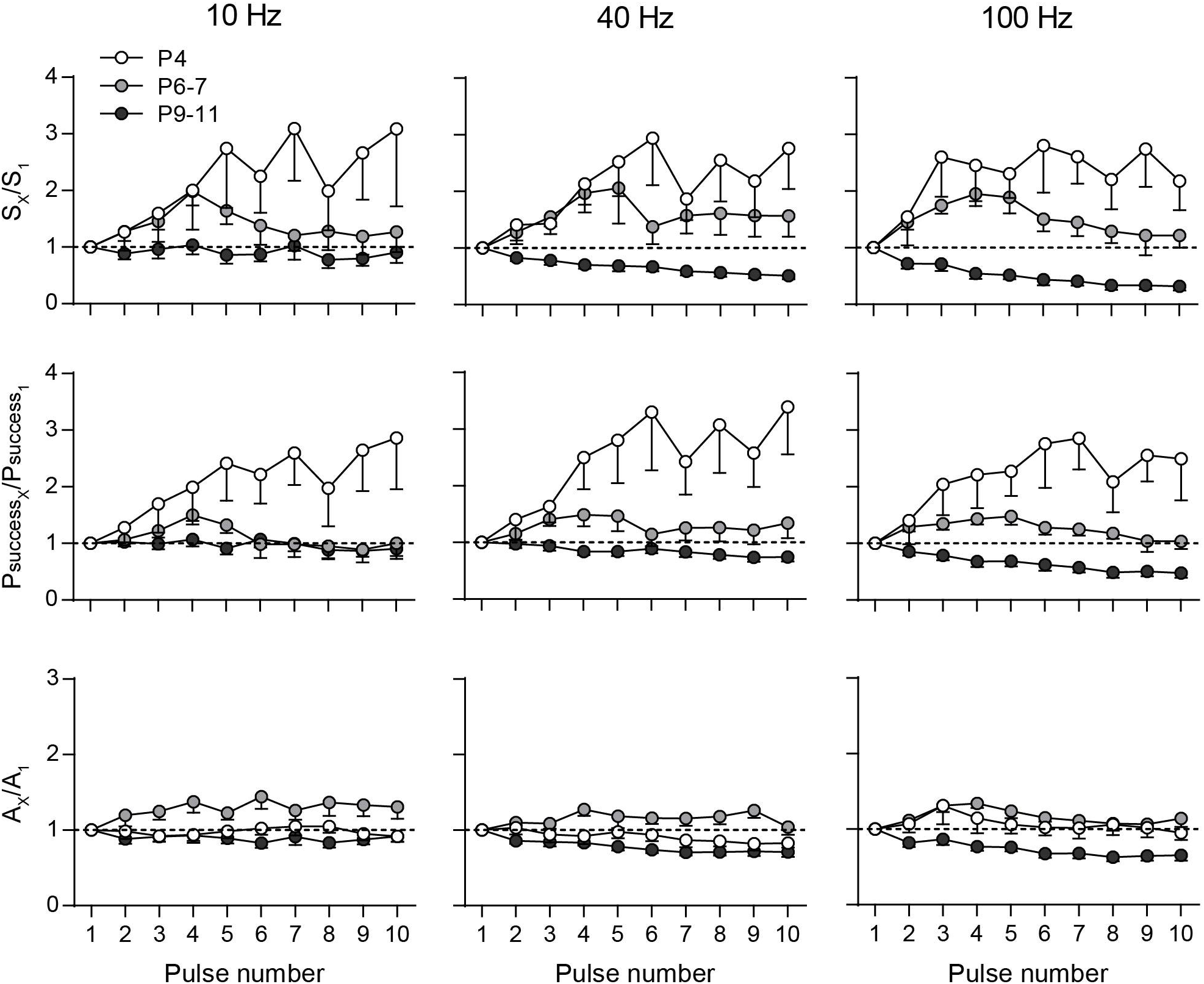
The STP pattern at the MOC-IHC synapse changes from facilitation to depression during development. The normalized overall response (S_X_/S_1_, upper panels), probability of successfully evoking a release event (P_successX_/P_success1_, middle panels) and average amplitude of successful responses (A_X_/A_1_, lower panels) were plotted against pulse number during 10-pulse stimulation trains at 10, 40 and 100 Hz. Experiments were done in IHCs at P4 (white circles), P6-7 (light grey circles) and P9-11 (dark grey circles). Error bars are S.E.M.

**Table 1.** Short-term synaptic plasticity parameters of the MOC-IHC synapse atdifferent stages of development. Stimulation trains consisting of 10 pulses at 10, 40 and 100 Hz were applied at P4, P6-7 and P9-11 IHCs. The overall response (S), the probability of successfully evoking a release event (P_success_), and the average amplitude of successful responses (A) were computed for each pulse. The first pulse values (X_1_) and the maximum (in case of facilitation) or minimum value (in case of depression) (X_max_(_min_)) for S, P_success_ and A are shown for each data set. *p* values correspond to one-way ANOVA/Friedman tests results when no significant differences were found, otherwise they correspond to the multiple comparisons post-hoc test result for X_max(min)_ shown. S and A values are in pA.

#### STP is sensitive to changes in the initial probability of release

Most synapses possess multiple forms of plasticity and the net synaptic strength depends on the interaction between them. Short term depression and facilitation are both present at a given synapse, but the relative prominence of each one is controlled by the initial probability of release (Regehr, 2012). As the probability of release is highly sensitive to the external Ca^2+^ concentration (Dodge and Rahamimoff, 1967), we evaluated whether changing this parameter by manipulating [Ca^2+^]_o_ affected the STP pattern at P6-7 and P9-11. At P6-7, increasing [Ca^2+^]_o_ from 1.3 to 1.5 mM caused a significant increase in the quantum content of evoked release (Ca^2+^1.3 mM, *m* = 0.36 ± 0.06, Ca^2+^1.5 mM, *m* = 0.69 ± 0.14, n = 10 cells, 6 mice, *p* = 0.0180, Figure 4 a). Under this condition, applying a 100 Hz 10 pulse-train, the STP pattern changed from facilitation to depression (Ca^2+^ 1.3 mM, n = 15 cells, 12 mice, Ca^2+^ 1.5 mM, n = 9 cells, 6 mice, Figure 4 b). Using the same rationale, at P9-11 we reduced [Ca^2+^]_o_ from 1.3 to 1.1 mM. This caused a significant reduction in the quantum content of evoked release (Ca^2+^ 1.3 mM, *m* = 1.43 ± 0.36, Ca^2+^1.1 mM, *m* = 0.89 ± 0.25, n = 7 cells, 5 mice, *p* = 0.0304, Figure 4 c) and a concomitant change in the STP pattern which changed from depression to facilitation (Ca^2+^ 1.3 mM, n = 16 cells, 12 mice, Ca^2+^ 1.1 mM, n = 10 cells, 6 mice, Figure 4 d).

**Figure 4.**
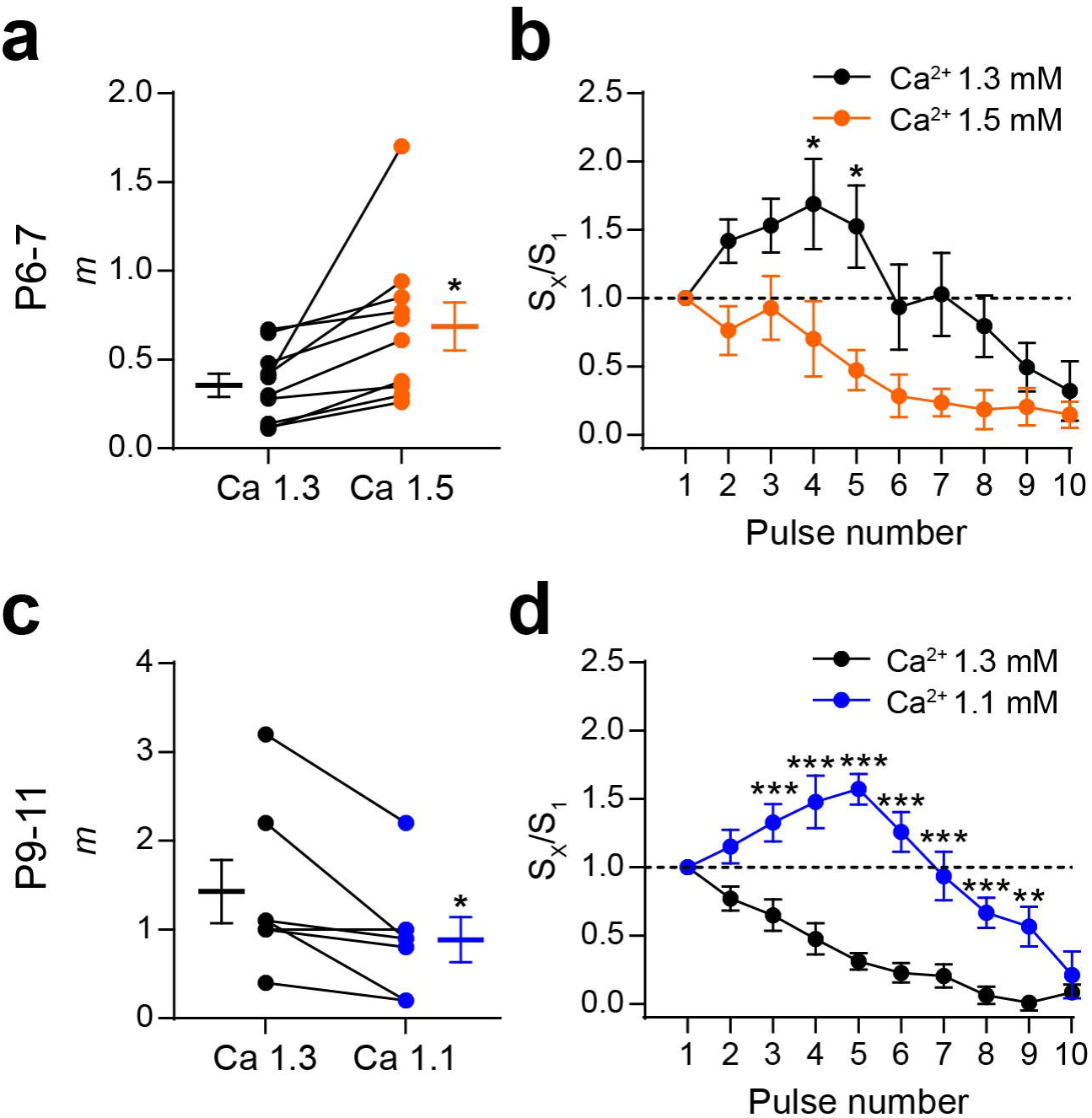
Changes in the initial probability of release modify the STP pattern. a, Graph showing that in P6-7 IHCs an increase in [Ca^2+^]_o_ from 1.3 (black circles) to 1.5 mM (orange circles) significantly increased m. The mean ± S.E.M. values of *m* measured in both [Ca^2+^]_o_ are plotted to the left and to the right of their respective individual responses. b, Normalized amplitude of eIPSCs (S_x_/S_1_) plotted against pulse number during 10-pulse stimulation trains at 100 Hz recorded in both [Ca^2+^]_o_ conditions in P6-7 IHCs. Note that the STP pattern changed from facilitation in [Ca^2+^]_0_ = 1.3 mM to depression in [Ca^2+^]_0_ = 1.5 mM. c, Graph illustrating the decrease in *m* at P9-11 IHCs after changing [Ca^2+^]_o_ from 1.3 to 1.1 mM. d, Normalized amplitude of eIPSCs (S_x_/S_1_) plotted against pulse number during 10-pulse stimulations trains at 100 Hz recorded in both [Ca^2+^]_o_ conditions in P9-11 IHCs. In this case, the STP pattern changed from depression at [Ca^2+^]_o_ = 1.3 mM to facilitation at [Ca^2+^] _o_= 1.1 mM. Error bars are S.E.M. \**p*<0.05, \*\**p*<0.01, \*\*\**p*<0.001.

#### Both the size of the readily releasable pool of vesicles and its replenishment rate increase during development

The increase in *m* observed during development can be due to changes either in the probability of release, in the quantum size or in the available number of synaptic vesicles (readily releasable pool, RRP). As reported above (Figure 2), no changes were observed in the quantum size during development, but there was both an increase in the probability of successfully evoking a release event (Figure 1) and also a significant increment in the frequency of spontaneous events across the three developmental stages studied. In order to test whether there were also changes in the RRP size at the MOC synaptic terminals contacting the IHCs during postnatal development, we applied high frequency stimulation trains (50 pulses at 100 Hz; 10-30 times) at each of the developmental stages studied (P4, P6-7 and P9-11). Representative examples of these trains are illustrated in Figure 5 (a, b and c). The cumulative mean amplitude along the train was calculated for each of the stimulation pulses and the last 20 points of the plot were fitted with a linear function. Representative cumulative amplitude plots are shown in Figure 5 (d, e and f). The intercept of the linear fit divided by the quantum size (mean amplitudes of sIPCs at each stage, Figure 2 c) is the RRP size and the slope of this linear regression is the rate at which the synapse replenishes the docking sites with new synaptic vesicles (Schneggenburger *et al.,* 1999; Neher, 2015). There was a significant increase in the size of the RRP between P4 and P9-11: P4 = 1.04 ± 0.28 vesicles; at P6-7 = 3.82 ± 0.81 vesicles and at P9-11 = 6.46 ± 1.12 vesicles (n = 12 cells, 10 mice, 14 cells, 9 mice and 15 cells, 7 mice for P4, P6-7 and P9-11, respectively). Significant differences were observed when comparing P4 vs P6-7 (*p* = 0.0093) and P4 vs P9-11 (p = 2.5e^−05^), but not P6-7 vs P9-11 (p = 0.1023; Figure 5 g, h). A significant increase of the replenishment rate between P4 and P11 was also observed. Replenishment rate at P4: 5.71 ± 1.39 vesicles/sec, at P6-7: 13.48 ± 2.81 vesicles/sec and at P9-11: 22.95 ± 3.33 vesicles/sec. Significant differences were observed when comparing P4 vs P6-7 (*p* = 0.034), P4 vs P9-11 (*p* = 2.4e^−05^) and P6-7 vs P9-11 (*p* = 0.032; Figure 5 i).

**Figure 5.**
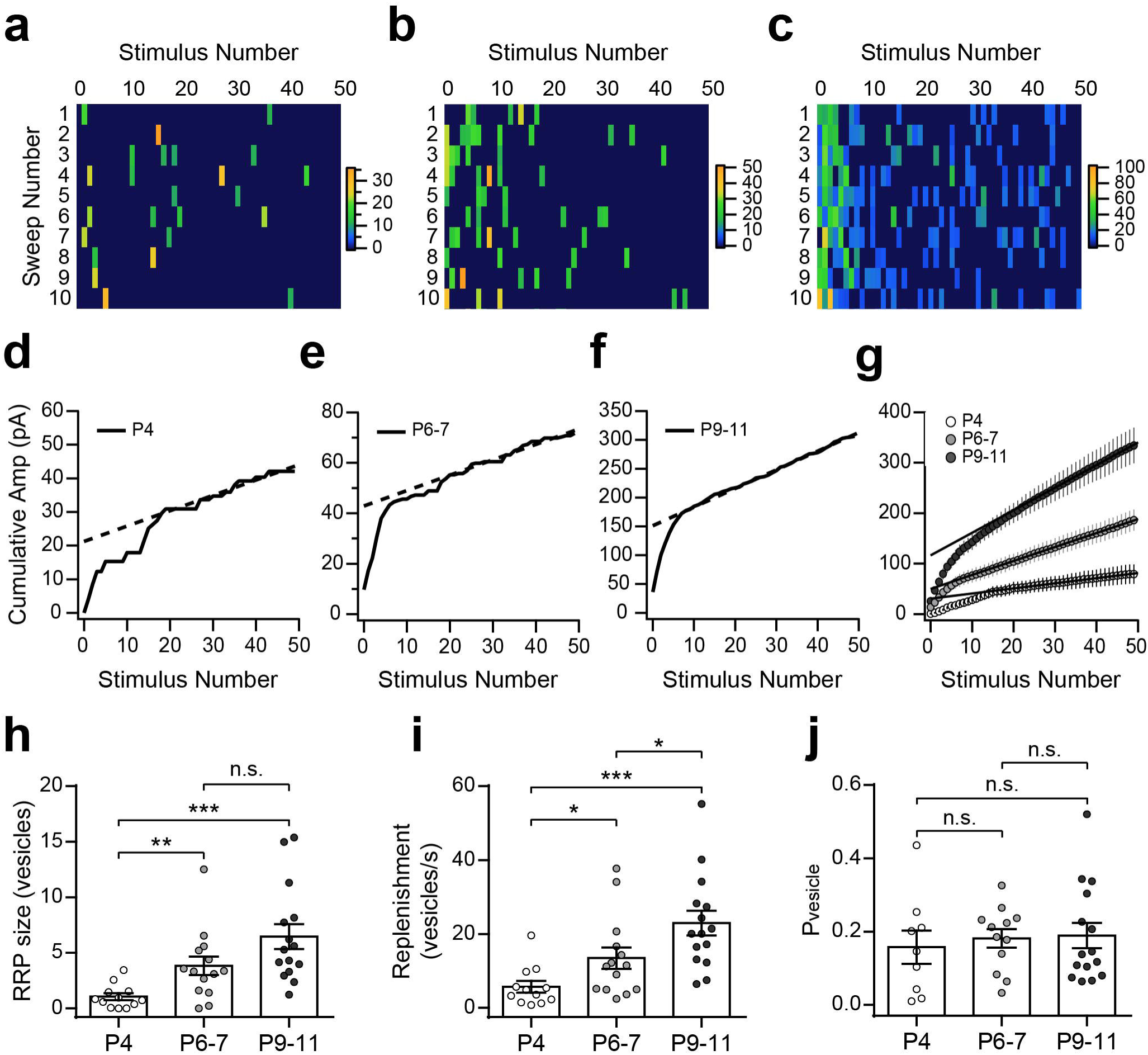
The size of the RRP and its replenishment rate increase during development. a, b and c, Color coded rectangles represent the amplitudes of the eIPSCs evoked by trains (50 pulses at 100 Hz, each panel shows 10 sweeps/columns) applied to P4, P6-7 and P9-11 IHCs. Color scale bars are expressed in pA. d, e and f, Representative examples of the average cumulative amplitude plots built to estimate the RRP and replenishment rate (continuous lines are the cumulative amplitudes and the dashed lines are the linear fits of the last 20 points of the plot). g, The cumulative plots built with the pooled data show the increase of the intercept at each of the developmental stages (P4: white; P6-7: light grey; P9-11: dark grey). h, Summary plot showing the increase in RRP size during development. I, Summary plot showing the increase in synaptic vesicles replenishment rate during high frequency stimulation at the same time period. j, Summary plot illustrating that P_vesicle_ remains constant throughout the developmental stages analyzed. Error bars are S.E.M. \**p*<0.05, \*\**p*<0.01, \*\*\**p*<0.001.

From these experimental data we estimated the probability of release of each vesicle, P_vesicle_ (Schneggenburger *et al.,* 1999, Valera *et al.,* 2012). By calculating the ratio between the average number of vesicles released per action potential (quantum content) and the number of vesicles ready to be released (RRP size), P_vesicle_ was obtained for each cell in which the RRP size was determined. No significant differences were observed in this parameter across the three stages studied (P4: P_vesicle_ = 0.16 ± 0.05, n = 9 cells, 8 mice; P6-7: P_vesicle_ = 0.18 ± 0.03, n = 12 cells, 7 mice; P9-11: P_vesicle_ = 0.19 ± 0.03, n = 15 cells, 7 mice; *p* = 0.825; Figure 5 j). Taken together, these results suggest that the increase in the RRP size and replenishment rate account for the increment in synaptic strength observed during development.

### Ion channels coupled to the release process at the transient MOC-IHC synapse during postnatal development

#### Developmental changes in Ca^2+^ channels coupled to ACh release

Ca^2+^ channels coupled to transmitter release have been shown to be developmentally regulated both at central synapses (Iwasaki *et al.,* 2000, Momiyama, 2003, Fedchyshyn and Wang, 2005) and at the NMJ (Rosato Siri and Uchitel, 1999). In a previous paper we showed, by electrophysiological and pharmacological methods, that transmitter release at the transient MOC-IHC synapse at P9-11 is supported by P/Q- and N-type VGCCs (Zorrilla de San Martín *et al.,* 2010). In view of the significant presynaptic functional changes found in synaptic strength and STP pattern between the different ages studied, we investigated whether the VGCC coupled to the release process at the transient MOC-IHC synapse were also developmentally regulated. We evaluated the quantum content of transmitter release at P4, P6-7 and P9-11 in the presence of toxins that specifically antagonize P/Q-, N- and R-type VGCC (Mintz and Bean, 1993, Olivera *et al.,* 1994, Newcomb *et al.,* 1998, Bourinet *et al.,* 2001). Omega-Agatoxin IVA (ω-Aga, 200 nM), ω-Conotoxin GVIA (ω-CgTx, 500 nM) and SNX-482 (SNX, 500 nM) were used to block P/Q-, N- and R-type VGCCs, respectively. The effects of these Ca^2+^ channel blockers on evoked release at the three postnatal stages studied are illustrated in Figure 6. At P4 and P6-7, both ω-Aga and SNX significantly reduced the quantum content of evoked release (P4: *m*_control_ = 0.27 ± 0.05, *m*_ω-Aga_ = 0.08 ± 0.03, n = 5 cells, 5 mice, *p* = 0.0029; m_control_ = 0.26 ± 0.07, *m*_SNX_ = 0.10 ± 0.03, n = 5 cells, 5 mice, *p* = 0.0295. P6-7: *m*_control_ = 0.44 ± 0.07, *m*_ω-Aga_= 0.17 ± 0.04, n = 5 cells, 5 mice, *p* = 0.0035; *m*_control_ = 1.22 ± 0.32, m_SNX_ = 0.50 ± 0.13, n = 5 cells, 5 mice, *p* = 0.0294). The N-type VGCC blocker (ω-CgTx), however, had no significant effect on evoked release at these two stages (P4: *m*_control_ = 0.46 ± 0.25, *m*_ω.CgTx_ = 0.40 ± 0.13, n = 3 cells, 3 mice, *p* = 0.6557. P6-7: *m*_control_ = 0.46 ± 0.09, *m*_ω-CgTx_ = 0.44 ± 0.05, n = 5 cells, 5 mice, *p* = 0.597). As previously reported, at P9-11, (Zorrilla de San Martín *et al.,* 2010), both the P/Q and the N-type channel blockers significantly reduced evoked release (note: data for these two toxins at this stage were taken from those used in (Zorrilla de San Martín *et al.,* 2010); they were tested for normal distribution, re-analyzed and illustrated for comparison). Thus, *m*_control_ = 0.86 ± 0.31, *m*_ω-Aga_ = 0.30 ± 0.07, n = 6 cells, 6 mice, *p* = 0.0313; *m*_control_ = 0.62 ± 0.12, *m*_ω-CgTx_= 0.35 ± 0.16, n = 5 cells, 5 mice, *p* = 0.014). At P9-11, the R-type VGCC blocker SNX did not have a significant effect on the quantum content of evoked release (*m*_control_ = 0.42 ± 0.14, *m*_SNX_ = 0.25 ± 0.07, n = 6 cells, 6 mice, *p* = 0.1563). Taken together, these results clearly show that there is a change in the VGCC subtypes coupled to the ACh release process during the critical period during which the MOC-IHC synapse is functional. At the earliest periods (P4 to 7), N-type VGCCs do not participate in evoked ACh release, which is supported by both P/Q and R-type VGCCs. At P9-11, both P/Q and N-type VGCC support release whereas at this stage R-type channels are not coupled to this process.

**Figure 6.**
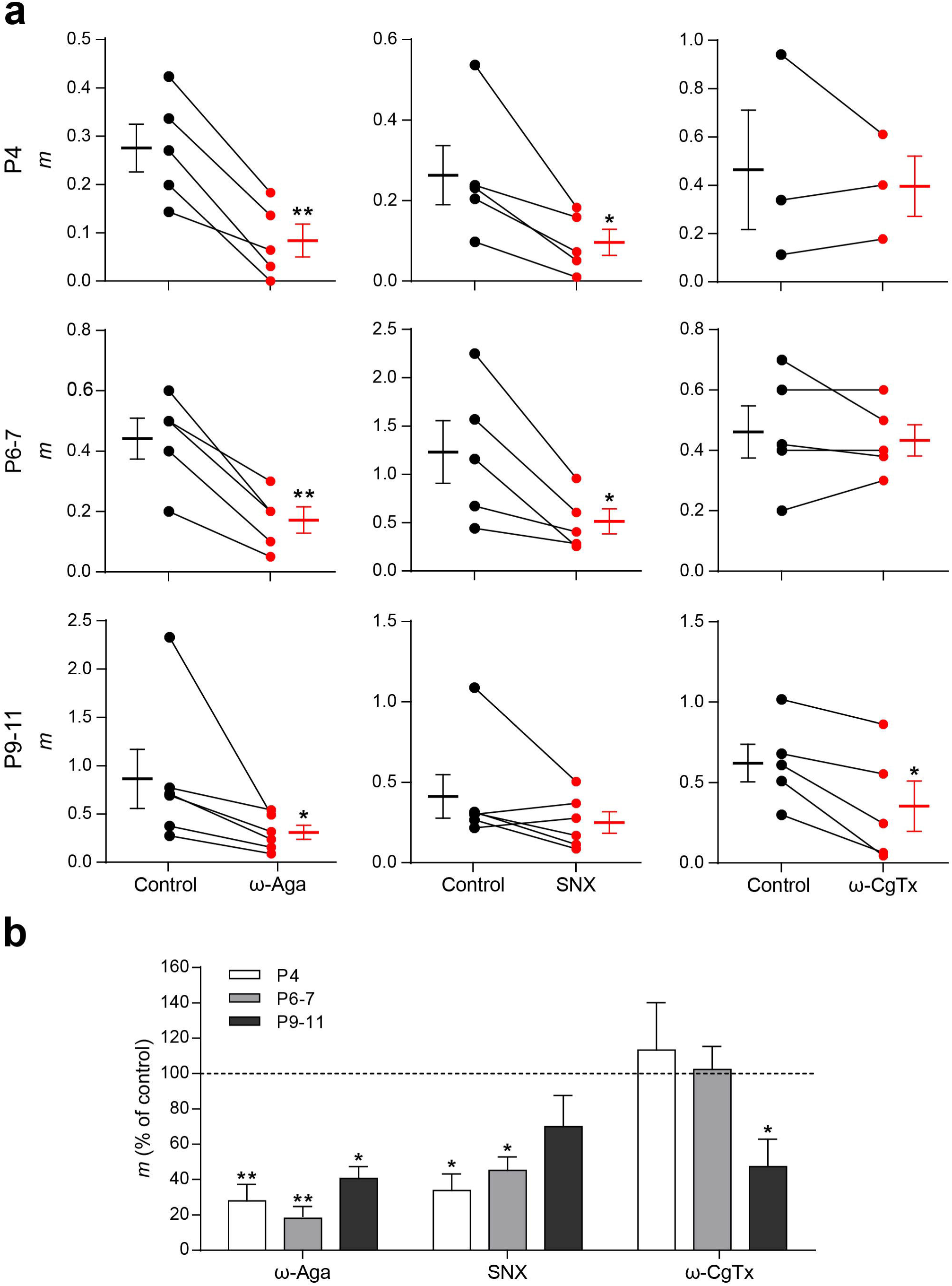
Different types of VGCCs support ACh release at the MOC-IHC synapseduring postnatal development. a, Graph showing the effect on *m* of 200 nM ω-Agatoxin IVA (ω-Aga), 500 nM SNX-482 (SNX) and 500 nM ω-Conotoxin GVIA (ω-CgTx), P/Q-, R- and N-type VGCCs specific antagonists, respectively, in P4, P6-7 and P9-11 cochlear preparations (control: black circles, treated: red circles). The mean ± S.E.M. values of *m* measured before and after toxin treatment are plotted to the left and to the right of their respective individual responses. b, Bar graph summarizing the effect on *m* of 200 nM ω-Aga, 500 nM SNX and 500 nM ω-CgTx at the different stages of development. Note that R-type VGCCs contribute to the release process only during early stages of development (P4 and P6-7), whereas N-type VGCCs begin to support ACh release at a later period (P9-11) (ω-Aga *m* % of control: P4 = 27.97 ± 9.30, P6-7 = 36.67 ± 6.48, P9-11 = 40.67 ± 6.79; SNX *m* % of control: P4 = 33.74 ± 9.39, P6-7 = 45.31 ± 7.61, P9-11 = 69.98 ± 17.72; ω-CgTx *m* % of control: P4 = 113.50 ± 26.79, P6-7 = 102.40 ± 13.00, P9-11 = 47.46 ± 15.47). Data from P9-11 mice are shown for comparison and were taken from Zorrilla de San Martín et al., 2010. Error bars are S.E.M. \**p*<0.05, \*\**p*<0.01.

### The role of L-type VGCC in the release process changes during development

L-type VGCCs do not support transmitter release at fast synapses under normal conditions (Catterall, 2011). However, they have been shown to participate in this process at reinnervating (Katz *et al.,* 1996) and developing neuromuscular synapses (Sugiura and Ko, 1997, Rosato Siri and Uchitel, 1999). It has also been reported that motoneurons possess silent L-type channels that may be recruited to facilitate transmitter release during high-frequency bursts (Oliveira *et al.,* 2004). In addition, L-type VGCCs are involved in the regulation of transmitter release at several synapses by the activation of Ca^2+^-dependent K^+^ conductances (Robitaille *et al.,* 1993, Robitaille *et al.,* 1993, Marrion and Tavalin, 1998, Prakriya and Lingle, 1999, Sun *et al.,* 2003, Loane *et al.,* 2007, Marcantoni *et al.,* 2007, Muller *et al.,* 2007, Berkefeld and Fakler, 2008, Fakler and Adelman, 2008, Grimes *et al.,* 2009). L-type VGCCs are highly sensitive to micromolar concentrations of dihydropyridines (DHP) that, either negatively (i. e., nifedipine, nitrendipine) or positively (i. e., (±) Bay-K 8644), modulate their activity (Doering and Zamponi, 2003, Catterall and Few, 2008). We have previously shown that at P9-11, incubation of the cochlear preparation with the L-type antagonist nifedipine (3 μM) causes a significant increase in the quantum content of evoked release, whereas incubation with the agonist Bay-K 8644 (10 μM) significantly reduces this parameter. This contradictory result was accounted for by the fact that at this stage, L-type channels are not directly involved in release but they are functionally coupled to the activation of Ca^2+^-activated BK channels that accelerate repolarization of the synaptic terminal during an action potential (Zorrilla de San Martín *et al.,* 2010). Therefore, the block of L-type VGCCs increased release whereas enhancement of their activity by Bay-K caused the opposite effect.

In order to study the role of L-type VGCCs in ACh release at the transient efferent synapse during development we tested the effects of nifedipine and Bay-K on *m* at P4 and P6-7. At P4 and P6-7, both the agonist (10 μM Bay-K) and the antagonist (3 μM nifedipine) of L-type VGCCs caused a significant increase in the quantum content of evoked release (Figure 7; P4: *m*_control_ = 0.12 ± 0.04, *m*_Bay-K_ = 0.35 ± 0.04, n = 6 cells, 6 animals, *p* = 0.0313; *m*_control_ = 0.29 ± 0.08, *m*_nife_ = 0.51 ± 0.12, n = 8 cells, 8 animals, *p* = 0.0078; P6-7: *m*_control_ = 0.73 ± 0.13, *m*_Bay-K_ = 1.74 ± 0.23, n = 6 cells, 6 animals, *p* = 0.0018; *m*_control_ = 0.72 ± 0.15, *m*_nife_ = 1.70 ± 0.27, n = 6 cells, 6 animals, *p* = 0.0043). As shown before (Zorrilla de San Martín *et al.,* 2010), at P9-11 (see also lower panel in Figure 7) nifedipine increased the quantum content of evoked release whereas Bay-K reduced it (*m*_control_ = 1.40 ± 0.12, *m*_Bay-K_ = 0.73 ± 0.14, n = 4 cells, 4 animals, *p* = 9.36e^−05^; *m*_control_ = 1.54 ± 0.19, *m*_nife_ = 3.20 ± 0.70, n = 7 cells, 7 animals, *p* = 0.02007). The fact that both the L-type agonist and antagonist increased release from P4 to P7 was surprising and led us to evaluate whether BK channels were also involved in modulating the release of ACh at these two earlier stages of development.

**Figure 7.**
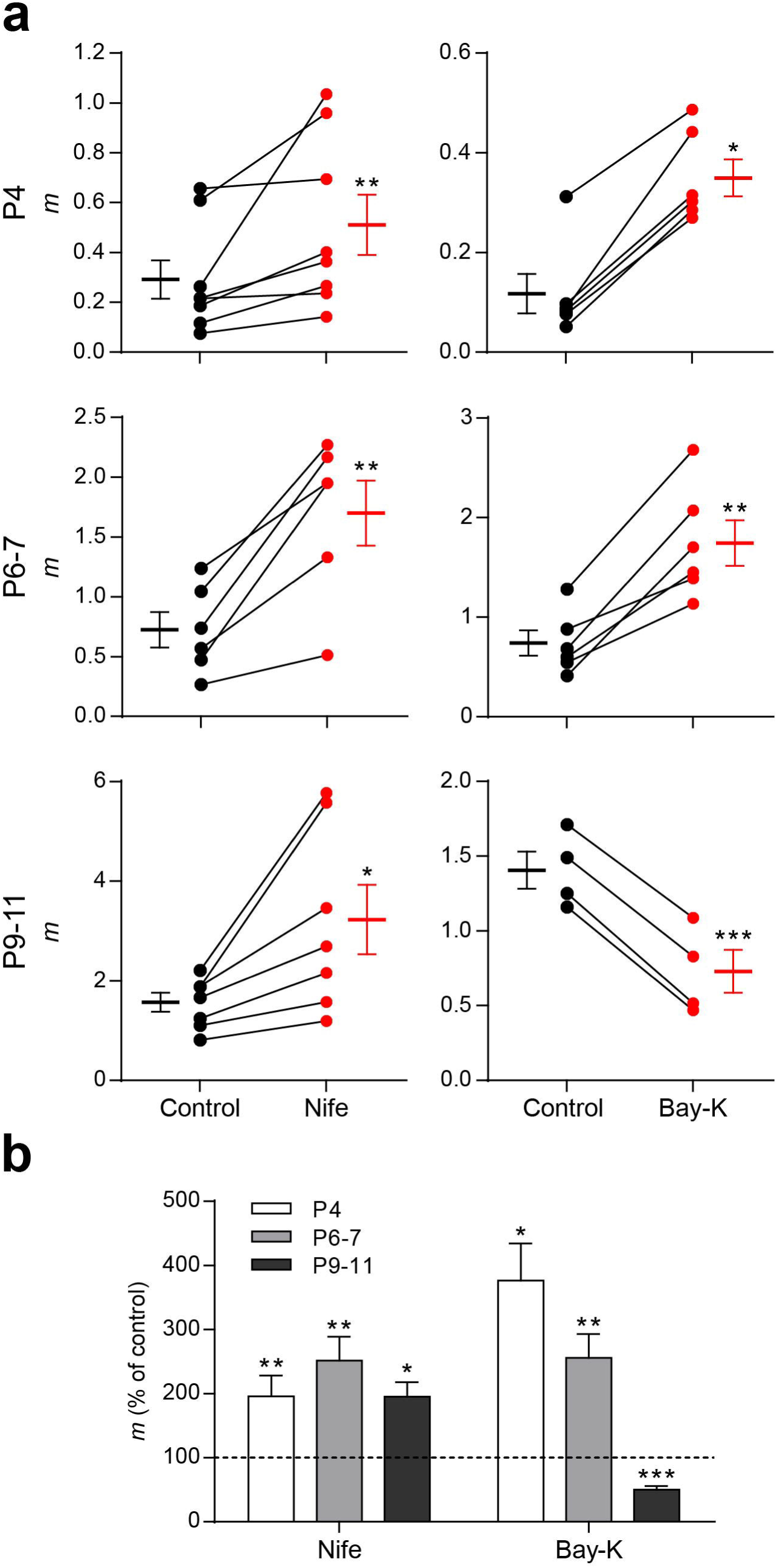
The role of L-type VGCC in transmitter release at the MOC-IHC synapsechanges during development. a, Graph showing the effect on *m* of 3 μM Nifedipine (Nife) and 10 μM Bay-K 8644, an antagonist and an agonist of L-type VGCCs, respectively, in cochlear preparations from mice at P4, P6-7 and P9-11 (control: black circles, treated: red circles). The mean ± S.E.M. values of *m* measured before and after toxin treatment are plotted to the left and to the right of their respective individual responses. b, Bar graph summarizing the effect on *m* of 3 μM Nife and 10 μM Bay-K at the different stages of development evaluated. At P4 and P6-7 mice both L-type VGCC antagonist and agonist increase *m* (Nife *m* % of control: P4 = 195.60 ± 32.45, P6-7 = 251.40 ± 37.24, P9-11 = 195.20 ± 22.58; Bay-K *m* % of control: P4 = 376.30 ± 58.11, P6-7 = 255.90 ± 37.25, P9-11 = 50.21 ± 5.61). Data from P9-11 mice are shown for comparison and were taken from Zorrilla de San Martín et al., 2010. Error bars are S.E.M. \**p*<0.05, \*\**p*<0.01, \*\*\**p*<0.001.

### BK channels are functionally expressed at the MOC-OHC synapse from P4 to P11

The participation of BK channels in the release process at the transient MOC-IHC synapse was evaluated by incubating the cochlear preparation with iberiotoxin (IbTx), a specific blocker of BK channels (Galvez *et al.,* 1990). Figure 8 illustrates de effects of 100 nM IbTx on the quantum content of ACh release at P4 (a, d), P6-7 (b, d) and P9-11 (c, d. data from Zorrilla de San Martín et al (2010) were tested for normal distribution, re-analyzed and plotted for comparison with the earlier stages). Blocking BK channels with IbTx caused a strong and significant increase in evoked release at all the developmental stages studied: P4: *m*_control_ = 0.12 ± 0.03, *m*_IbTx_ = 0.25 ± 0.04, n = 12 cells, 12 animals, *p* = 0.0068. P6-7: *m*_control_ = 1.34 ± 0.19, *m*_IbTx_ = 2.65 ± 0.44, n = 8 cells, 8 animals, *p* = 0.0018. P9-11: *m*_control_= 1.57 ± 0.37, *m*_IbTx_ = 3.88 ± 0.77, n = 6 cells, 6 animals, *p* = 0.0313. This result indicates that BK channels negatively modulate the release of ACh at the three stages studied.

**Figure 8.**
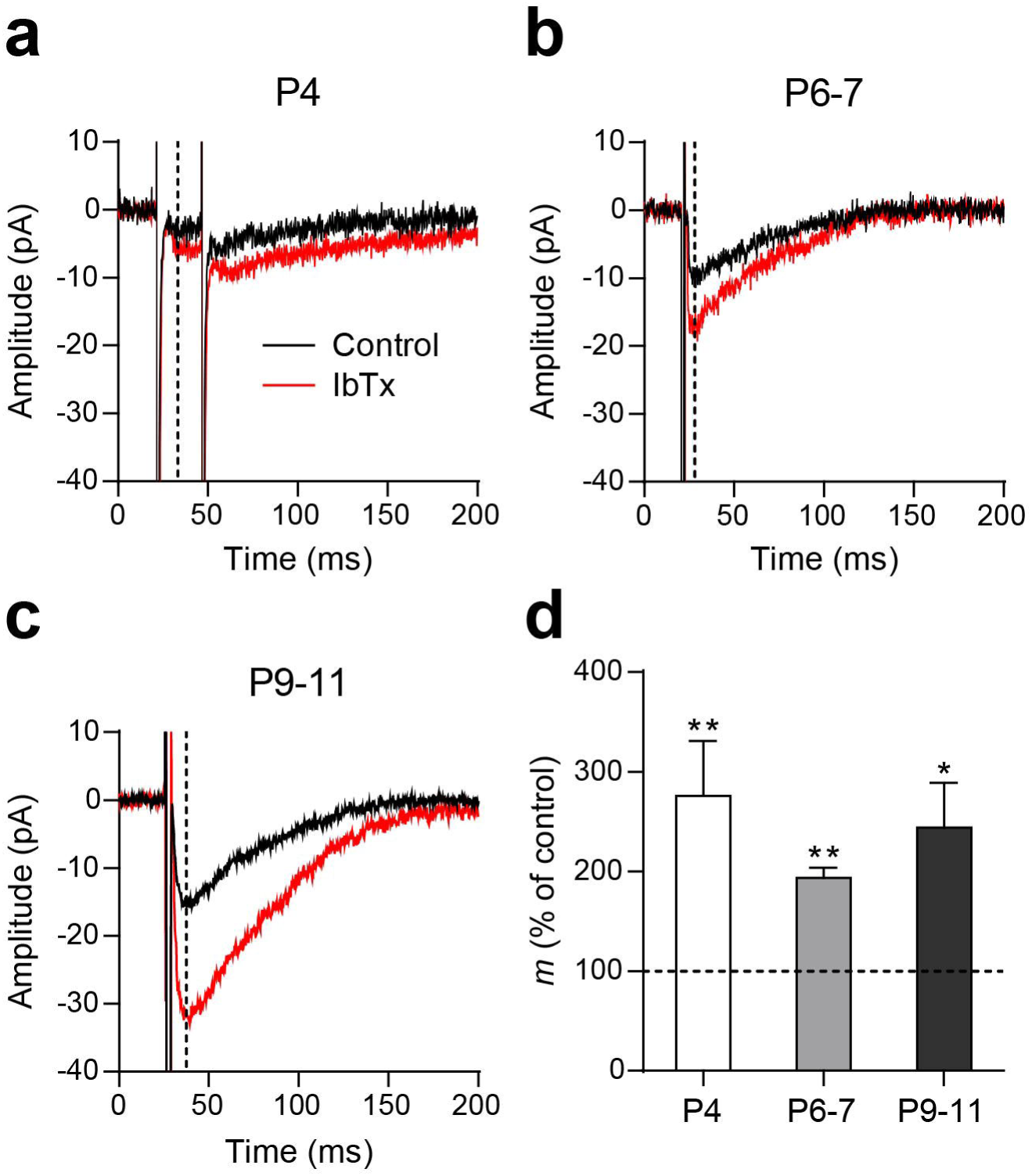
BK channels negatively modulate ACh release at the MOC-IHC synapse throughout development. a, b and c, Graphs showing the effect on *m* of 100 nM Iberiotoxin (IbTx), an antagonist of BK channels, at P4 (a), P6-7 (b) and P9-11 (c) cochlear preparations (control: black circles, treated: red circles). The mean ± S.E.M. values of *m* measured before and after toxin treatment are plotted to the left and to the right of their respective individual responses. b, Bar graph summarizing the effect on *m* of 100 nM IbTx at the different stages of development evaluated (IbTx *m* % of control: P4 = 275.50 ± 55.05, P6-7 = 196.80 ± 10.01, P9-11 = 263.30 ± 47.62). Data from P9-11 mice are shown for comparison and were taken from Zorrilla de San Martín et al., 2010. Error bars are S.E.M. \**p*<0.05, \*\**p*<0.01.

### Ca^2+^ influx through L-type VGCC plays a dual role at early developmental stages

We have previously shown that at P9-11, preincubation of the cochlear preparation with the BK channel blocker IbTx (100 nM) completely occludes the effects of both nifedipine and Bay-K (Zorrilla de San Martín *et al.,* 2010). Therefore, we now tested the effects of Bay-K on the quantum content of evoked release in cochlear preparations from P4 and P6-7 mice preincubated with IbTx (100 nM). Contrary to that observed at P9-11, even in the absence of functional BK channels, Bay-K significantly increased *m* at both P4 and P6-7 MOC-IHC synapses (P4: *m*_Control_ = 0.25 ± 0.19, *m*_IbTx_ = 0.45 ± 0.22, *m*_IbTx+Bay-K_ = 0.79 ± 0.34, n = 5 cells, 5 animals, IbTx vs control *p* = 0.0021, IbTx+Bay-K vs IbTx *p* = 0.0021, IbTx+Bay-K vs control *p* = 1.9e^−05^; P6-7: *m*_Control_ = 1.16 ± 0.28, *m*_IbTx_ = 2.27 ± 0.67, *m*_IbTx+Bay-K_ = 3.31 ± 0.78, n = 5 cells, 5 animals, IbTx vs control *p* = 0.0248, IbTx+Bay-K vs IbTx *p* = 0.041, IbTx+Bay-K vs control *p* = 9.88e^−07^; P9-11: *m*_Control_ = 1.19 ± 0.29, *m*_IbTx_ = 2.63 ± 0.82, *m*_IbTx+Bay-K_ = 2.54 ± 0.63, n = 6 cells, 6 animals, IbTx vs control *p* = 0.0107, IbTx+Bay-K vs IbTx *p* = 0.9809, IbTx+Bay-K vs control *p* = 0.0189, Figure 9 a).

**Figure 9.**
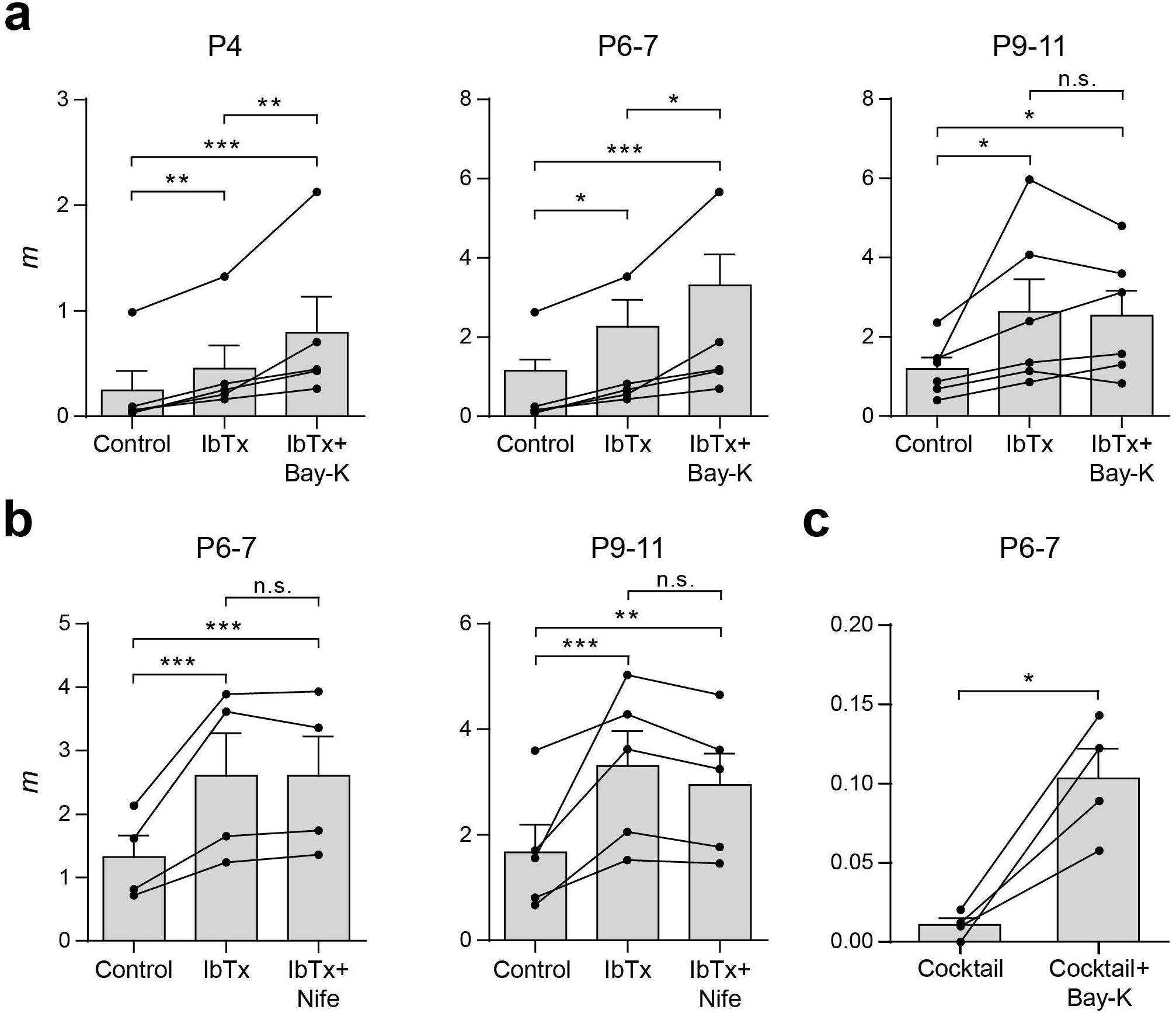
L-type VGCCs play a dual role on ACh release at the earlier stages of development. a, Summary plots showing that 10 μM Bay-K significantly increased *m* after blocking BK channels with 100 nM IbTx only in P4 and P6-7 IHCs. Values from each individual cell are shown as black circles. b, Summary plots illustrating that 3 μM nifedipine did not further increase *m* after preincubating the cochlear preparations with 100 nM IbTx at P6-7, as previously observed at P9-11. c, Summary plot showing that 10 μM Bay-K was still able to increase *m* after blocking P/Q-, R- and N-type VGCCs and BK channels with a toxin cocktail containing 200 nM ω-Aga, 500 nM ω-CgTx, 500 nM SNX and 100 nM IbTx. Data from P9-11 mice are shown for comparison and were taken from Zorrilla de San Martín et al., 2010. Error bars are S.E.M. \**p*<0.05, \*\**p*<0.01, \*\*\**p*<0.001.

As the effects of DHPs on evoked release were similar at P4 and P6-7 and given the much lower number of functionally innervated IHCs found at P4, the following experiments were carried out only at P6-7. Surprisingly, nifedipine (3 μM) had no effect on release at P6-7 after having blocked BK channels (*m*_Control_ = 1.32 ± 0.34, *m*_IbTx_ = 2.60 ± 0.67, *m*_IbTx+Nife_ = 2.60 ± 0.62, n = 4 cells, 4 animals, IbTx vs control *p* = 6.82e^−06^, IbTx+Nife vs IbTx *p* = 1, IbTx+Nife vs control *p* = 6.86e^−06^, Figure 9 b). This lack of effect of the L-type VGCC antagonist was the same as that observed at P9-11 (*m*_control_ = 1.67 ± 0.52, *m*_IbTx_ = 3.30 ± 0.66, *m*_IbTx+Nife_ = 2.95 ± 0.59, n = 5 cells, 5 animals, control vs IbTx *p* = 3.47e^−04^, IbTx vs IbTx+Nife *p* = 0.6789, control vs IbTx+Nife *p* = 0.0073, plotted here for comparison (Zorrilla de San Martín *et al.,* 2010). The effect of Bay-K suggests that at the earlier stages L-type VGCCs are involved in supporting release. However, the lack of effect of nifedipine suggests the contrary.

In view of these conflicting results between the effects of Bay-K and nifedipine in the absence of functional BK channels (Figure 9 a and b), we decided to confirm the participation of L-type VGCCs in transmitter release by testing the effects of Bay-K in the presence of P/Q-, R- and N-type VGCC and BK channel blockers (the ion channel blocking cocktail was made up of 200 nM ω-Aga, 500 nM ω-CgTx, 500 nM SNX and 100 nM IbTx). After preincubating the cochlear preparation with this cocktail, Bay-K significantly increased *m* (*m*_cocktail_ = 0.011 ± 0.004, *m*_cocktail+Bay-K_ = 0.103 ± 0.019, n = 4 cells, 4 mice, *p* = 0.0152, Figure 9 c). This confirms that at P6-7, under certain conditions, Ca^2+^ flowing in through L-type VGCCs can reach the evoked-release site machinery and support ACh release.

At P9-11 BK channels are activated only by Ca^2+^ influx through L-type VGCC without the participation of Ca^2+^ flowing in through either P/Q or N-type VGCCs (Zorrilla de San Martín *et al.,* 2010). Therefore, we also tested whether this was also the case at P6-7 by evaluating the effects of IbTx after preincubating the cochlear preparation with nifedipine (3 μM). As illustrated in Figure 10 (P6-7: *m*_control_ = 0.64 ± 0.21, *m*_Nife_ = 1.44 ± 0.34, m_Nife+IbTx_ = 1.70 ± 0.39, n = 4 cells, 4 animals, control vs Nife *p* = 0.0039, Nife vs Nife+IbTx *p* = 0.8622, control vs Nife+IbTx *p* = 5.66e^−05^; P9-11: *m*_control_ = 1.34 ± 0.13, m_Nife_ = 2.53 ± 0.35, *m*_Nife+IbTx_ = 2.45 ± 0.47, n = 5 cells, 5 animals, control vs Nife *p* = 0.0174, Nife vs Nife+IbTx *p* = 0.9809, control vs Nife+IbTx *p* = 0.0294), nifedipine also occluded the effects of IbTx at P6-7 (P9-11 values are from Zorrilla de San Martín *et al,* 2010, and plotted for comparison). This shows that throughout development BK channels at MOC-IHC synapses are only activated by Ca^2+^ influx through L-type VGCCs.

**Figure 10.**
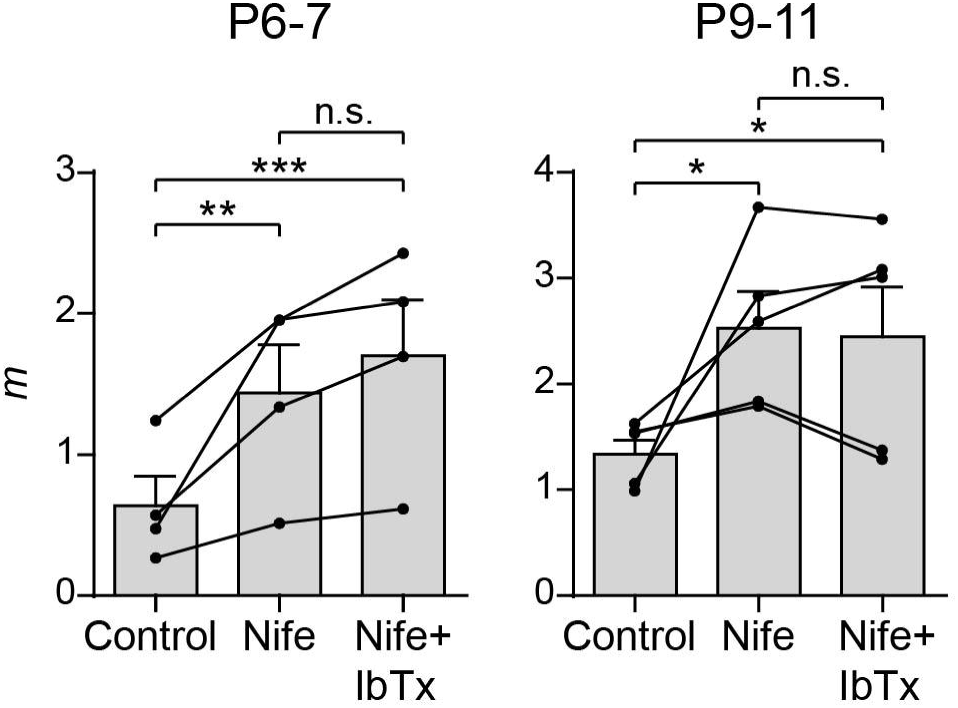
BK channels are activated only by Ca influx through L-type VGCCs. Summary plots showing that 100 nM IbTx did not further increase *m* after preincubating cochlear preparations with 3 μM Nife at P6-7 (left panel), as previously described for P9-11 (right panel, Zorrilla de San Martín et al., 2010). Values from each individual cell are shown as black circles. Error bars are SEM. \**p*<0.05, \*\**p*<0.01, \*\*\**p*<0.001.

### Effects of L-type channel modulators on K^+^-evoked release

In order to further investigate whether the differential effects of the DHPs on electrically evoked ACh release during development could be related to changes in the co-localization of L-type VGCCs with respect to the release sites activated upon invasion of an action potential to the synaptic terminal, we tested the effects of Bay-K and nifedipine on K^+^-evoked release of transmitter. This type of release is independent from the generation of an action potential due to Na+ channel inactivation under constant depolarization (Hodgkin and Huxley, 1952). Therefore, K^+^-evoked release is independent from the effects of BK channel activation on the repolarization of the terminal action potential.

Figure 11 illustrates the effects of Bay-K (10 μM) and nifedipine (3 μM) at P6-7 (upper panels in a and b) and at P9-11 (lower panels in a and b). In this type of release the L-type agonist Bay-K significantly increased the frequency of sIPSCs at both ages (P6-7 sIPSC frequency control 0.80 ± 0.23 Hz, sIPSC frequency Bay-K 5.65 ± 1.56 Hz, n = 6 cells, 6 animals, *p* = 0.0205; P9-11 sIPSC frequency control 2.25 ± 1.19 Hz, sIPSC frequency Bay-K 5.98 ± 1.93 Hz, n = 7 cells, 7 animals, *p* = 0.0313) while the antagonist nifedipine significantly reduced it (P6-7 sIPSC frequency control 2.69 ± 0.90 Hz, sIPSC frequency Nife 0.61 ± 0.11 Hz, n = 7 cells, 7 animals, *p* = 0.0156; P9-11 sIPSC frequency control 3.42 ± 1.74 Hz, sIPSC frequency Nife 1.40 ± 0.70 Hz, n = 6 cells, 6 animals, *p* = 0.0313). This result suggests that the differences found in the effects of Bay-K on P4 and P6-7 with respect to P9-11 might be related to developmental changes in the localization of L-type channels with respect to the action potential evoked-neurotransmitter release machinery.

**Figure 11.**
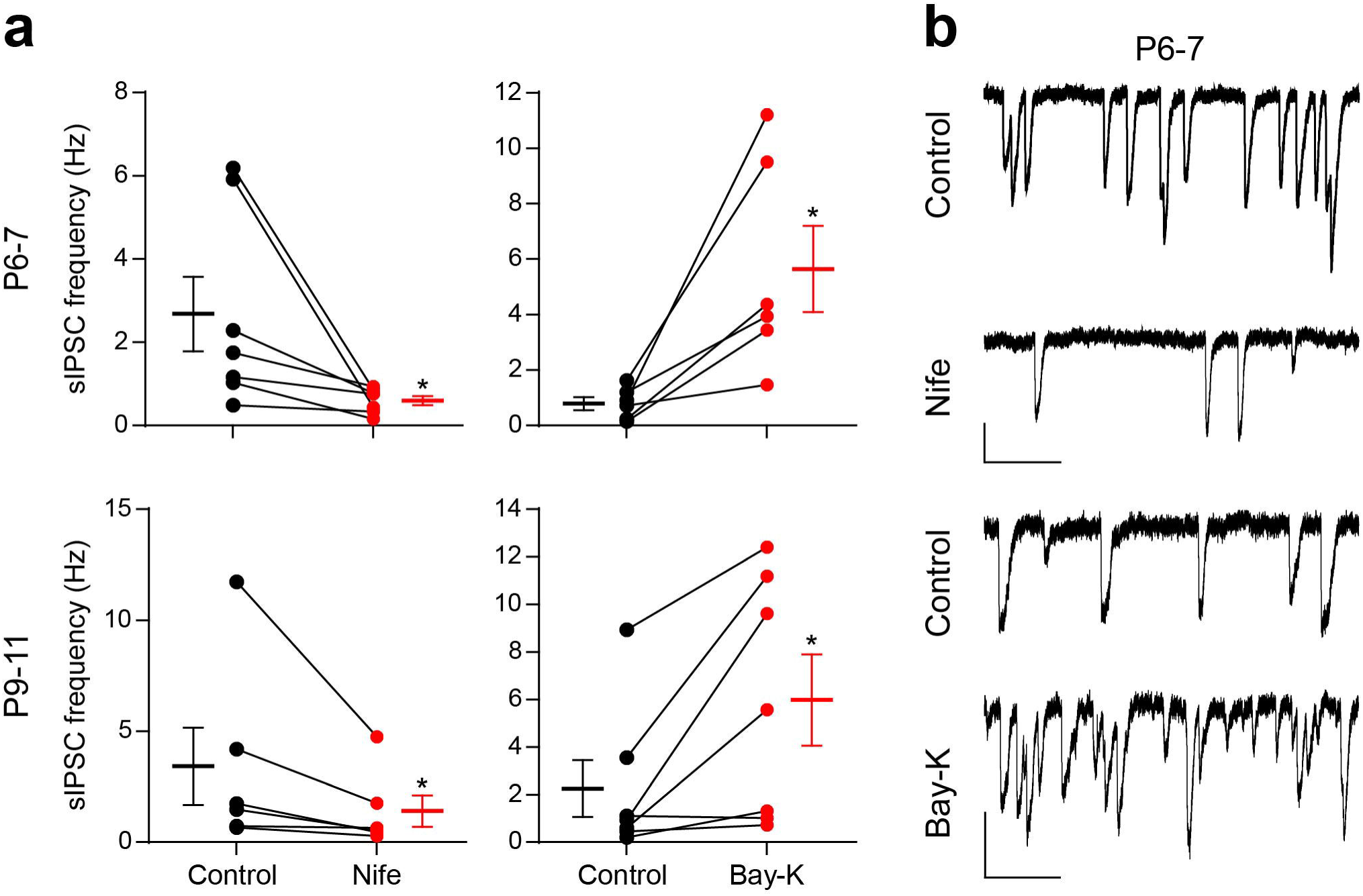
Effect of DHPs on transmitter release evoked by high external potassium. a, Graphs showing the effect of 3 μM Nifedipine (Nife) (left panels) and 10 μM Bay-K 8644 (right panels) on the frequency of sIPSC recorded at P6-7 IHCs (upper panels) and P9-11 IHCs (lower panels) during application of an external solution containing 15-25 mM K^+^. b, Representative traces recorded before and after incubation with 3 μM Nife (upper panels) and 10 μM Bay-K (lower panels) at P6-7 IHCs. Vertical scale bars: 50 pA, horizontal scale bars: 1 s. Error bars are S.E.M. \**p*<0.05.

## Discussion

### Developmental changes in the biophysical properties of ACh release

The MOC-IHC synapse is functional from P0 to hearing onset. Both the number of functionally innervated cells and the amplitude of ACh-evoked currents significantly increase from P3 to P11 returning to very low values just after hearing onset. In addition, the α10 nAChR subunit and the SK2 channel, two key postsynaptic proteins, are down-regulated and disappear at ∼P14 (Katz *et al.,* 2004, Roux *et al.,* 2011). We now show that during the developmental critical period before hearing onset, dramatic presynaptic changes take place at the transient MOC-IHC synapse.

Synaptic strength significantly increased from P4 to P9-11 due to a 5.6x increase in the quantum content with a concomitant reduction in the number of release failures. This, together with the increase in sIPSCs frequency and the lack of changes in their amplitude and kinetics, support the notion that the increment in synaptic strength is of presynaptic origin (Del Castillo and Katz, 1954). The fact that sIPSC amplitude remained constant indicates that neither the quantum size nor the postsynaptic sensitivity to ACh changed from P4 to P11. The sIPSC decay time constant is dominated by the closure kinetics of SK2 channels (Oliver *et al.,* 2000), therefore as already reported (Katz *et al.,* 2004, Roux *et al.,* 2011), coupling between the α9α10 nAChR and the SK2 channel is stable from P4 to P11.

Changes in quantum content were reflected in the STP pattern which shifted from facilitation at P4 and P6-7 to depression at P9-11. Variations in STP during development have been observed in other systems, namely, the visual cortex (Chen and Roper, 2004), cerebellum (Pouzat and Hestrin, 1997), hippocampus (Schiess *et al.,* 2010) and the calyx of Held (Taschenberger and von Gersdorff, 2000). Curiously, in the rat MOC-IHC synapse at P9-11, facilitation of responses is observed upon high frequency stimulation (Goutman *et al.,* 2005), whereas in mice we found depression under the same experimental condition. As the quantum content at both synapses is similar (∼1), other factors account for these species differences in STP pattern. This could include the RRP size or replenishment rate, the VGCCs supporting and/or modulating release, differential modulation by BK channels (Zorrilla de San Martín *et al.,* 2010), nitric oxide (Kong *et al.,* 2013) and other neurotransmitter systems (Wedemeyer *et al.,* 2013, Ye *et al.,* 2017).

There is accumulated evidence indicating that facilitation is caused by an elevated [Ca^2+^]_i_ remaining from the previous stimulus (Jackman and Regehr, 2017). The relationship between the increment in the quantum content and short-term depression is usually accounted for by a greater RRP depletion due to a greater number of neurotransmitter quanta released at the first stimulus of the train (Zucker and Regehr, 2002). These notions are consistent with our results showing that changes in the quantum content modified the STP pattern: at P9-11 depression upon 100 Hz stimulation train was replaced by facilitation upon reducing [Ca^2+^]_o_ from 1.3 to 1.1 mM, whereas at P6-7 facilitation was replaced by depression upon raising it from 1.3 to 1.5 mM. During development, however, P_vesicle_ remained constant whereas RRP and its replenishment rate increased. Therefore, RRP depletion is not likely to be the cause of depression upon high frequency stimulation. It has been reported that currents through P/Q-type VGCCs can either facilitate or depress upon repetitive stimulation depending on the presence of regulatory proteins (Catterall *et al.,* 2013, Nanou and Catterall, 2018). Therefore, differences in the expression of regulatory proteins might account for the changes in STP pattern between P9-11 and earlier developmental stages (Rosato Siri and Uchitel, 1999, Wu *et al.,* 1999). The amount of transmitter release evoked by an action potential depends upon the size of the effective RRP and on the initial probability of release of each of the vesicles within this pool (Thanawala and Regehr, 2013). Therefore, as P_vesicle_ remained constant, the increment observed in P_success_ is likely to be accounted for by the increment in RRP size. This agrees with the observation that quantum content and RRP size across the three ages studied presented a similar incremental ratio (P6-7/P4: *m* = 3, RRP size = 3.6; P9-11/P6-7: m = 2, RRP size = 1.7; P9-11/P4: *m* = 5.6, RRP size = 6.2).

### Developmental changes in voltage-gated Ca^2+^ channels coupled to ACh release

We have shown that at P9-11 transmitter release at the MOC-IHC synapse is mediated by Ca^2+^ influx through P/Q- and N-type VGCCs (Zorrilla de San Martín *et al.,* 2010). We now show that P/Q-type VGCCs are coupled to ACh release from P4 to P11. However, at P4 and P6-7, N-type VGCCs do not participate in release which is mediated by P/Q- and R-type VGCCs. In addition, at P9-11 there is a switch from R-type to N-type VGCCs. The participation of R-type VGCCs in ACh release from MOC fibers is in agreement with PCR analysis in the mouse cochlea (Green *et al.,* 1996) and with immunohistochemical studies which show that Ca_v_2.3 (R-type) is expressed by MOC fibers from P2 to P14 (Waka *et al.,* 2003). In addition, Ca_v_2.1 (P/Q-type) has been also shown to be expressed in the mammalian inner ear by transcriptome analysis (Gabashvili *et al.,* 2007).

Developmental changes in VGCCs coupled to transmitter release have been reported both at the NMJ (Rosato Siri and Uchitel, 1999) and at central synapses (Iwasaki *et al.,* 2000, Momiyama, 2003, Fedchyshyn and Wang, 2005). At those synapses, both P/Q and N-type VGCCs are coupled to the release process at early stages of maturation. In adults, P/Q-type channels are the predominant VGCCs coupled to transmitter release. The contribution of N-type VGCCs either completely disappears during maturation in the NMJ (Rosato Siri and Uchitel, 1999), in GABAergic synapses of the cerebellum and thalamus (Iwasaki *et al.,* 2000) and glutamatergic synapses of the calyx of Held (Fedchyshyn and Wang, 2005) or significantly diminishes in GABAergic striatal synapses (Momiyama, 2003). Curiously, at the MOC-IHC synapse N-type VGCCs do not mediate transmitter release at the earliest developmental stages.

At immature neuromuscular and calyx of Held synapses P/Q-type VGCCs are closer to the calcium sensor than N-type VGCCs and therefore it has been suggested that this spatial distribution accounts for the greater efficacy of P/Q-type channels in coupling to release (Rosato Siri and Uchitel, 1999, Wu *et al.,* 1999). In this respect, the observation that P_vesicle_ did not change with maturation of the MOC-IHC synapse might be related to the fact that P/Q-type channels mediate this process across all the developmental stages studied. Therefore, at the MOC-IHC synapse P/Q-type channels might be closer to the release sites compared to the other VGCCs that also support this process and thus be the main regulators of release probability.

### The dual role of L-type VGGCs at early stages of development

Based on the effects of L-type channel modulators and the BK channel antagonist on transmitter release at the MOC-IHC synapse, we show that at the early stages of development Ca^2+^ influx through L-type VGCCs has a dual role. On the one hand, it exerts a negative feedback on transmitter release as it activates BK channels which accelerate terminal membrane repolarization (Storm, 1987, Berkefeld and Fakler, 2008), thereby reducing the amount of transmitter released per action potential (Robitaille *et al.,* 1993, Robitaille *et al.,* 1993, Raffaelli *et al.,* 2004, Zorrilla de San Martín *et al.,* 2010). On the other hand, under certain conditions, as observed in the presence of Bay-K which increases the mean open time of L-type VGCCs (Bechem and Hoffmann, 1993) or when BK channels and all other VGGCs are blocked, Ca^2+^ influx through L-type VGCCs can reach the release sites and contribute to ACh release.

The dual role of VGCCs on transmitter release has also been demonstrated at the frog NMJ for N-type VGCCs where the entry of Ca^2+^ that triggers release also activates BK channels (Robitaille *et al.,* 1993, Robitaille *et al.,* 1993). The fact that as of P4 to P7, Ca^2+^ influx through L-type VGGCs, apart from activating BK channels, can also reach the release sites whereas at P9-11 it cannot, strongly suggests that at the early stages there is less compartmentalization of the presynaptic proteins that compose the action potential evoked-release machinery. Even though other experimental approaches should be employed to give further support to this hypothesis at the MOC-IHC synapse, spatial tightening during development between the proteins involved in the release process has been described both at the calyx of Held synapse (Fedchyshyn and Wang, 2005, Leao and von Gersdorff, 2009) and at cerebellar cortical synapses (Baur *et al.,* 2015).

## Conclusions

Variation in the expression and localization of ion channels involved in synaptic transmission at the MOC-IHC synapse together with changes in synaptic strength (summarized in Figure 12) underlie developmental changes in STP. Moreover, changes in the STP pattern during the short life of the MOC-IHC synapse might lead to fine tuning of IHC action potential frequency pattern (Goutman *et al.,* 2005, Johnson *et al.,* 2011, Sendin *et al.,* 2014), thus regulating signaling at the first synapse of the auditory pathway during its establishment.

**Figure 12.**
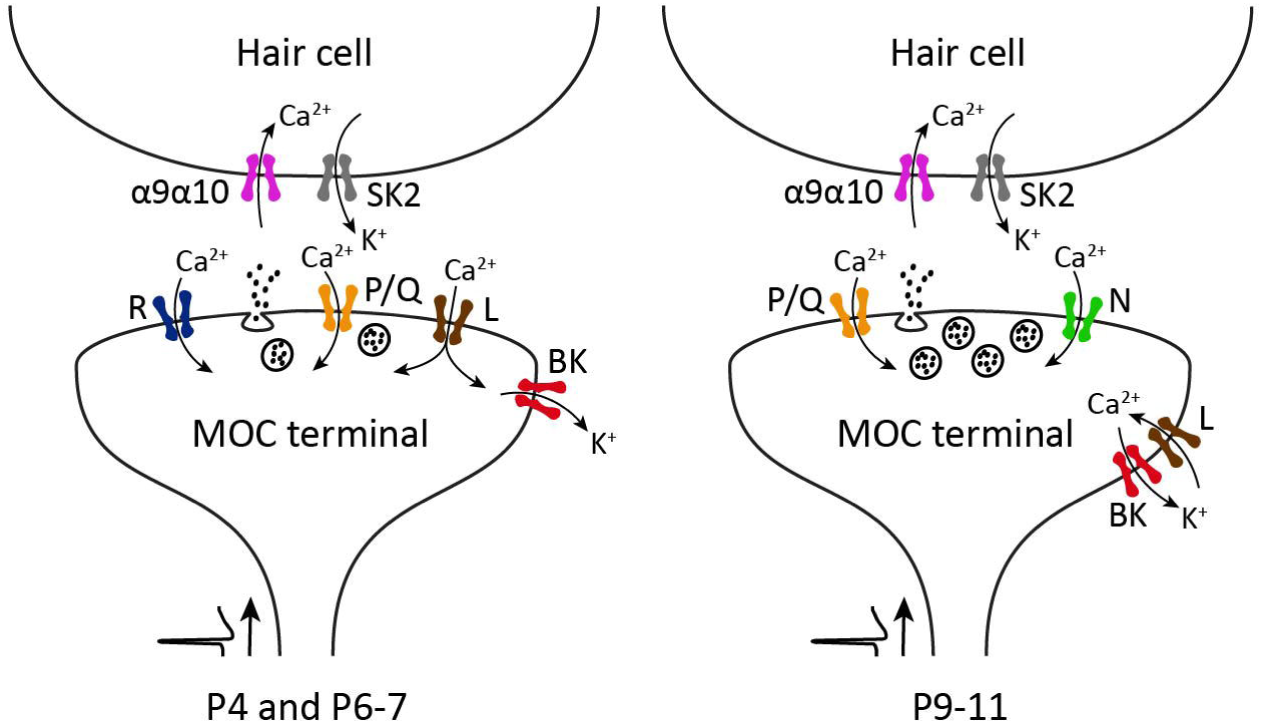
Schematic representation of the ion channels that support and modulate ACh release from MOC presynaptic terminals transiently innervating the IHCs during postnatal development. After an action potential arrives to the presynaptic terminal, Ca^2+^ influx through P/Q-, R- and L-type VGCCs mediate ACh release at the earlier stages of development (P4 and P6-7). At a later stage (P9-11), R-type VGCCs are down-regulated and N-type VGCC start contributing to the release process. Throughout this period, Ca^2+^ entering through L-type VGCCs exert a negative modulation on transmitter release by activating BK channels, which accelerate the repolarization of the terminal action potential thereby reducing the amount of neurotransmitter released per nerve impulse.

## Acknowledgements

This work was supported by Agencia Nacional de Promotion Cientifica y Tecnologica, Argentina (A.B.E. and E.K.), University of Buenos Aires, Argentina (E.K. and A.B.E.) and NIH Grant R01 DC001508 (Paul A. Fuchs, A.B.E.).

Javier Zorrilla de San Martín present address: Institut du Cerveau et de la Moelle épinière – ICM CNRS UMR 7225 - Inserm U1127 - UPMC-P6 UMR S 1127

The authors declare no competing financial interests

